# CRISPRa-mediated activation of Tfeb in a fibril-induced alpha-synuclein aggregation model

**DOI:** 10.64898/2026.10.16.757246

**Authors:** Elin Arvidsson, Elsa Pi Börjesson, Ermelinda Sabarese, Luis Quintino, Cecilia Lundberg

**Affiliations:** CNS Gene Therapy, Department of Experimental Medical Science, Lund University, Lund, Sweden; Danish Research Institute of Translational Neuroscience (DANDRITE), Nordic EMBL Partnership for Molecular Medicine and Department of Biomedicine, Aarhus University, Denmark

**Keywords:** Parkinson’s disease, Alpha-synuclein, TFEB, CRISPR-activation

## Abstract

Accumulation of misfolded alpha-synuclein (α-syn) is a key hallmark of Parkinson’s disease (PD), making it a key target for novel therapeutic strategies. Here, we use a previously described screening assay to select multiple efficient single guide RNA (sgRNA) combinations targeting transcription factor EB (TFEB) for CRISPR-activation (CRISPRa) using a nuclease-deficient *Staphylococcus aureus* Cas9 coupled to a truncated VP64-p53-Rta (VPR) activator complex. Selected sgRNA are applied in a rat neuroblastoma in-vitro model of fibril-induced α-syn aggregation with the goal of inducing autophagy and reduce protein aggregation. We successfully induce α-syn aggregation in this model and observe a stagnation in phosphorylated α-syn (p-α-syn) compared to non-phosphorylated α-syn across treatment groups. While we achieve robust expression of *Tfeb* in the exogenous screening assay and at endogenous mRNA level, we did not observe proportional increase in protein expression. Additionally, we do not observe any additive effects of CRISPRa induction combined with media treatments inducing Tfeb nuclear translocation. Taken together, these findings illustrate difficulties in epigenetic induction of Tfeb using CRISPRa and highlights the need for additional studies exploring this application.

## 1. Introduction

Parkinson’s disease (PD) is the second most prevalent neurodegenerative disease world-wide, currently estimated to affect 10 million people (1). The progressive nature of PD leads to progressive degradation of cell bodies located within the basal ganglia, deep subcortical nuclei with a central role in motor control (2). Specifically, the deterioration of the dopaminergic neurons found in the substantia nigra (SN) and the fibres connecting these cells to the striatum (STR), dubbed the nigro-striatal pathway, leads to classical parkinsonian signs, including rigidity and a loss of voluntary motor control (3). While PD has been extensively studied for decades and today’s treatment options can provide significant relief for patients, curative therapies have yet to be developed.

A key hallmark in PD pathology is the formation of heterogenous protein aggregates consisting primarily of alpha-synuclein (α-syn) (4, 5). Physiological α-syn is predominantly located presynaptically and has been shown to play key roles in regulation of vesicle trafficking and subsequent neurotransmitter release (6). Contrastingly, misfolded α-syn monomers and their subsequent formation of oligomers and fibrils have been suggested to contribute to the dysfunction and eventual demise of dopaminergic neurons through numerous mechanisms. This includes mitochondrial dysfunction, neuroinflammation, and inhibition of autophagy (7-9). The sensitivity of these cells to pathological α-syn highlights α-syn aggregation as an attractive target for new therapeutic developments.

Numerous methods have been developed for studying the effects of α-syn aggregation in both healthy and disease contexts, including a range of animal- and in vitro models (10-12). One strategy involves addition of synthetically produced pre-formed α-syn fibrils (PFFs) to cell cultures with or without overexpression of α-syn, resulting in formation of protein aggregates (13, 14). Using various cell culture approaches to model α-syn pathology provides a time- and cost-efficient way of investigating molecular mechanisms underlying disease, thereby carrying potential for multiplexing approaches and pharmacological screens.

Previous studies using α-syn aggregation models have suggested that impairment of autophagy may contribute to disease progression, and conversely, that increasing or restoring physiological autophagy response may prevent accumulation of neurotoxic aggregate species (9, 15). Transcription factor EB (TFEB) is a master transcriptional regulator of a broad range of genes related to autophagy and lysosomal biogenesis and has previously been suggested as target of interest in protein aggregation disorders, including PD (15, 16). Phosphorylated TFEB is retained in the cytoplasm during physiological conditions but is dephosphorylated and translocates to the nucleus in response to starvation or chemical induction (17, 18). While autophagy disruption contributes to cell death, excessive autophagic responses may have similarly detrimental effects on cell survival, thus stressing the importance of a delicate approach when manipulating this response (19, 20).

Despite growing interest in TFEB-based therapies, epigenetic activation of TFEB remains poorly characterized. CRISPR activation (CRISPRa) amplifies endogenous gene expression by utilizing an endonuclease-deficient (“dead”) Cas9 complex coupled to cis- or trans-activator elements, often utilizing multiple sgRNA to achieve optimal gene induction (21-23). Compared to transgenic knock-in or cDNA-mediated overexpression, the recruitment of endogenous activator domains through CRISPRa may achieve a more physiologically relevant level of gene expression.

Previous work by our group has described a streamlined workflow for cloning and screening one or multiple sgRNA candidates for downstream CRISPRa applications (24). This system utilizes a previously described miniaturized activator (Mini-dCas9v2), consisting of a truncated VP64-p53-Rta (VPR) activator complex combined with a nuclease-deficient *Staphylococcus aureus* Cas9 (Sa-dCas9).

Here, we employ this system to screen and select dual sgRNA candidates for targeted upregulation of endogenous *Tfeb* with the goal of suppressing PFF-induced α-syn aggregation in a rat glioblastoma cell line model of synucleinopathy.

## 2. Materials and methods

### Cloning and screening assay

Assembly of rat Tfeb sgRNA constructs was performed as previously described (23). Briefly, spacer sequences were generated using CHOP-CHOP (25) and designed as Gene Art Strings (Thermofisher) before being assembled in blunt end TOPO vectors (Thermofisher, #450245). TOPO vectors containing the respective sgRNA cassettes were co-transfected with a pBG.CMV.Mini-dCas9v2 vector containing a miniaturized, enzymatically dead Cas9 (dCas9) from *Staphylococcus aureus*; a reporter plasmid containing the target promoter sequence of interest coupled to TdTomato; and a transfection control plasmid containing a BFP cassette under control of an EF1a promoter. Transfection was performed in HEK-293T cells. Readout from the fluorescent reporter construct in the respective conditions was normalized against the BFP control to estimate efficiency of exogenous promoter activation between spacer sequences. Selected sgRNA spacer sequences were assembled in a single construct using Golden Gate cloning as previously described (24). The assembled sgRNA cassette was then excised and inserted in a pHG-Ef1a-mRFP acceptor vector through standard plasmid subcloning.

### Cell culture

C6 rat glioblastoma cells were purchased from the American Type Culture Collection (ATCC®, CCL-107). Cells were cultured in Ham’s F-12K (Kaighn’s) Medium (Gibco, #21127022) supplied with 10% FBS (Cytiva, #SV30160.03) and 1% Penicillin-streptomycin (Gibco, #15070063) at 37°C and 5% CO_2_.

### Plasmid transfection

C6 rat glioblastoma cells were seeded in a 6-well format at 300,000 cells per well and maintained as described above. Twenty-four hours after seeding, cultures were co-transfected with pBG.CMV.Mini-dCas9v2 and the respective guide combinations at a 2:1 dCas9:sgRNA plasmid ratio at a total DNA concentration of 600 ng. Forty-eight hours after transfection, RNA was prepared using the PureLink™ RNA Mini Kit (Invitrogen, #12183018A) according to manufacturer’s instructions. A total of 1 ug RNA was used for cDNA synthesis using iScript™ Reverse Transcription Supermix for RT-qPCR (Bio-Rad, #1708841) according to manufacturer’s instructions. The 20 ul cDNA reaction was diluted in 80 uL nuclease-free H^2^O for a final volume of 100 ul before RT-qPCR analysis.

### Viral vector production and transduction

HEK-293T cells (11×10^6^) were seeded in a T175 flask. Twenty-four hours later, cultures were transfected at a 3:1 PEI:DNA ratio, using 4,3 ug VSV-G-envelope plasmid (Addgene #12259), 8,5 ugp8.91 helper plasmid (Addgene #187441), and transgene for a total of 30 ug DNA per flask. Forty-eight hours post-transfection, cultures were harvested and centrifuged briefly at 800xg at 4°C. The supernatant was pushed through a 0.45 μm filter and centrifuged at 25,000 RPM, 4°C, for 90 minutes. The viral pellet was resuspended in DPBS and incubated at 4°C for 16 hours before aliquoting. For titration, 1,5e5 HEK-293T cells were transduced with serial dilutions of the viral suspension. Seventy-two hours later, total DNA was harvested using a DNeasy Blood & Tissue kit (QIAGEN, #69506), and RT-qPCR analysis was used to determine the viral titer using standard delta-delta-Ct calculations. Primers (Eurofins Genomics) and FAM-TAM hydrolysis probes (Roche, #04887301001) targeting human albumin (forward: 5’-TGA AAC ATA CGT TCC CAA AGA GTT-’3, reverse: 5’-CTC TCC TTC TCA GAA AGT GTG CAT AT-’3, probe: 5’-[FAM]-T GCT GAA ACA TTC ACC TTC CAT GCA GA-[TAM]-3’) and WPRE (forward: 5’-GGC ACT GAC AAT TCC GTG GT-’3, reverse: 5’-AGG GAC GTA GCA GAA GGA CG-3’, probe: 5’-[FAM]-A GCT CCT TTC CAT GGC TGC C-[TAM]-3’) were used.

### RNA extraction and cDNA preparation

C6 rat glioblastoma cells were seeded in a 12-well format at 40,000 cells per well and maintained as described above. Four hours after seeding, cells were co-transduced with pBG.CMV.Mini-dCas9v2 and respective sgRNA combinations at an MOI of 15 per construct. 72 hours after transduction, cells were collected and snap-frozen before being stored at -80°C until RNA extraction. RNA was prepared using the PureLink™ RNA Mini Kit (Invitrogen, #12183018A) according to manufacturer’s instructions. cDNA was prepared from a total of up to 1 ug RNA using iScript™ Reverse Transcription Supermix for RT-qPCR (Bio-Rad, #1708841) according to manufacturer’s instructions.

### Gene expression analysis

Relative gene expression was quantified as described previously (23). Briefly, RT-qPCR reactions were prepared at 10 uL total reaction volume containing 1 ul cDNA sample, 4 ul primer mix, and 5 ul SybrGreen mastermix. Analysis was performed in the LightCycler 480 Real-Time PCR instrument (Roche). Amplification was performed according to the following protocol: pre-incubation (95°C, 5 min), 45 cycles of denaturation (95°C, 10s), annealing (60°C, 10s), and extension (72°C, 10s). Melting curve was performed at 65°C, increasing in ramp rate at 4.8 °C/s up to 95°C with a hold time of 5 min. Values were analyzed using standard delta-delta-Ct calculations. The following primers were used: Tfeb (forward: AGAGCATGGCTCCCCTTTCC, reverse: AGCCAGGGTCACAGAACGTC) and actin (forward: TCTCTTCCAGCCTCC, reverse: TAGAGCCACCAATCC).

### PFF transfection

C6 cells were seeded in 12-well plates at a density of 20,000 cells per well. The next day, human pre-formed alpha synuclein fibrils (PFFs) produced in-house from human alpha synuclein monomers (MJFF, Proteos, # SKU: RP-010) or BSA were diluted to a working concentration of 400 ng/ul. For each well, 12 ul PFF- or BSA solution at working concentration and 63 ul Opti-MEM were prepared. Diluted PFFs were sonicated using a cuphorn sonicator at 70% amplitude with 1 second cycles and 1 second intervals, for 3 minutes. Sonicated samples were centrifuged briefly to avoid dispersion of droplets. For each sample, 6 ul of Lipofectamine 2000 reagent (Invitrogen, #11668-027) and 69 ul of Optimem (Gibco, #31-985-070) were prepared and incubated at RT for 5 minutes. The lipofectamine and PFF-solutions were mixed at a 1:1 ratio and incubated at RT for 10 minutes. A total of 125 ul of the final transfection mixture was added to each well. For 96-well plates, 10 ul was added to each well. Samples were incubated 24h before analysis.

### Immunocytochemistry

Ninety-six-well PhenoPlates (Revvity, #6055300) were coated with 100 ul poly-L-orthonine (PLO) at a concentration of 20 ug/ml and kept at 37°C overnight. Plates were washed three times with DPBS and dried for at least 2 hours before seeding. C6 cell lines were seeded at a confluency of 5000 cells per well and maintained as described above. Four hours after seeding, cultures were transduced at a MOI of 7,5 per construct. Seventy-two hours post-transduction, cultures were fixed in 1% PFA for 10 minutes at room temperature (RT) and washed three times with DPBS. Cultures were subsequently washed three times with KPBS before being incubated in 200 ul 5% blocking solution (5% serum from secondary antibody species and 0,25% TritonX in 1x KPBS) for one hour at RT. After blocking, cultures were incubated in 200 ul primary antibody suspension at 4°C overnight. The next day, cultures were washed three times in 1x KPBS, incubated in blocking solution for one hour at RT, and subsequently incubated with secondary antibody solution and respective dyes for one hour at RT. Cultures were washed three times with 1x KPBS before being conserved in 0,1% sodium azide diluted in 1x KPBS and stored at 4°C until analysis. Image analysis and fluorescence quantification were performed using the Operetta CLS system (Revvity). Antibodies and dyes were prepared as follows: rabbit-anti-TFEB (1:200, Protein Tech, #13372-1-AP), rabbit-anti-LC3B (1:200, Sigma, #L7543), rabbit-anti-LAMP1 (1:1000, Abcam, #ab24170), mouse-anti-alpha synuclein (1:1000, Santa Cruz, #sc-12767), rabbit-anti-alpha synuclein (phospho S129) (1:1000, Abcam, #ab51253), chicken anti-GFP (1:1000, Abcam, ##ab13970) goat-anti-rabbit Alexa fluor 647 (1:400, Invitrogen, #A-21244), donkey-anti-mouse Alexa 647 (1:500, Invitrogen, ##A-31571), goat-anti-chicken Alexa 488 (1:500, Invitrogen, #A-11039), phalloidin labeling probes (Invitrogen, #A30106), and DAPI (1:1000).

### FACS analysis (FloIT)

FACS aggregate analysis was performed based on the FloIT technique previously established by Whiten and colleagues (26). C6 cells were cultured in a 12-well plate as described in the method section above (“PFF transfection”). Twenty-four hours after transfection, the media was aspirated, and the wells were gently washed with 1 ml PBS before the addition of 150 ul 0,05% trypsin (Gibco, #11538876) and incubation at 37°C for 5 minutes. Detached cells were resuspended in 850 ul cell medium. A non-transfected control well was counted to estimate average cell density and remaining samples were resuspended to an estimated density of 150,000 cells/ml. Samples were centrifuged at 400xg for 5 min at RT, and resuspended in lysis buffer (1.5 mL/sample) containing Triton X-100 (5ul/ml), RedDot™2 Far-Red Nuclear Stain (0,5ul/ml) (Biotium, #40061), and cOmplete™ EDTA-free Protease Inhibitor Cocktail tablets (1 tablet/10mlsef1324) (Roche, # 04693159001) in PBS. Samples were incubated in lysis buffer at RT for two minutes and then placed on ice until analysis. Analysis was performed in the LSRFortessa (BD Biosciences) with wavelength acquisition settings for GFP (488/509 nm) and RedDot2 (622/694 nm). The following voltages were used: 350 (FSC), 220 (SSC), 479 (Alexa Fluor 700/RedDot2), and 423 (FITC/GFP). The nuclear fraction was identified based on co-expression of GFP and RedDot2 and extra-nuclear fraction based on expression of GFP. All samples were run at low speed to permit small protein detection. Import of FSC-files and subsequent analysis was performed in FlowJo version 10 (FlowJo).

### Statistics

Statistical analyses were performed in GraphPad Prism 10. Ordinary One-way ANOVA was performed to assess differences between group means. Equality of variance was assessed using the Brown-Forsythe test, and multiple comparisons were performed against a control group using Dunnett’s test (alpha =0.05). When assessing multi-variable group means, Two-Way ANOVA was performed using Tukey’s Honestly Significant Difference test and Dunnett’s test (alpha =0.05).

## 3. Results

### Assessment of selected sgRNA candidate combinations for Tfeb induction

To evaluate sgRNA candidates for targeted induction of the *Tfeb* promoter, we utilized an exogenous sgRNA screening assay previously developed in-house. Seven sgRNA spacer sequences targeting the region 600 bp upstream of *the Tfeb* transcription start-site (TSS) in *Rattus norvegicus* were generated. sgRNA were cloned as cassettes containing individual human U6 (hU6) promoters and transfected alone, or in combination, in HEK-293T cultures to assess activation potential. (Fig 1, Fig S1). Eight separate sgRNA combinations resulted in statistically significant reporter activation compared to a scrambled transfection control (Fig. 1B). Three sgRNA combinations with the highest estimated promoter activation efficiency—sgRNA 3+5, 3+6, and 3+7—were selected for downstream applications. Selected sgRNA cassettes including individual hU6 promoters were assembled in a single plasmid construct and co-delivered with Mini-dCas9V2 activator into C6 rat glioblastoma cell cultures through lentiviral (LV) transduction or chemical transfection.

**Figure 1.**
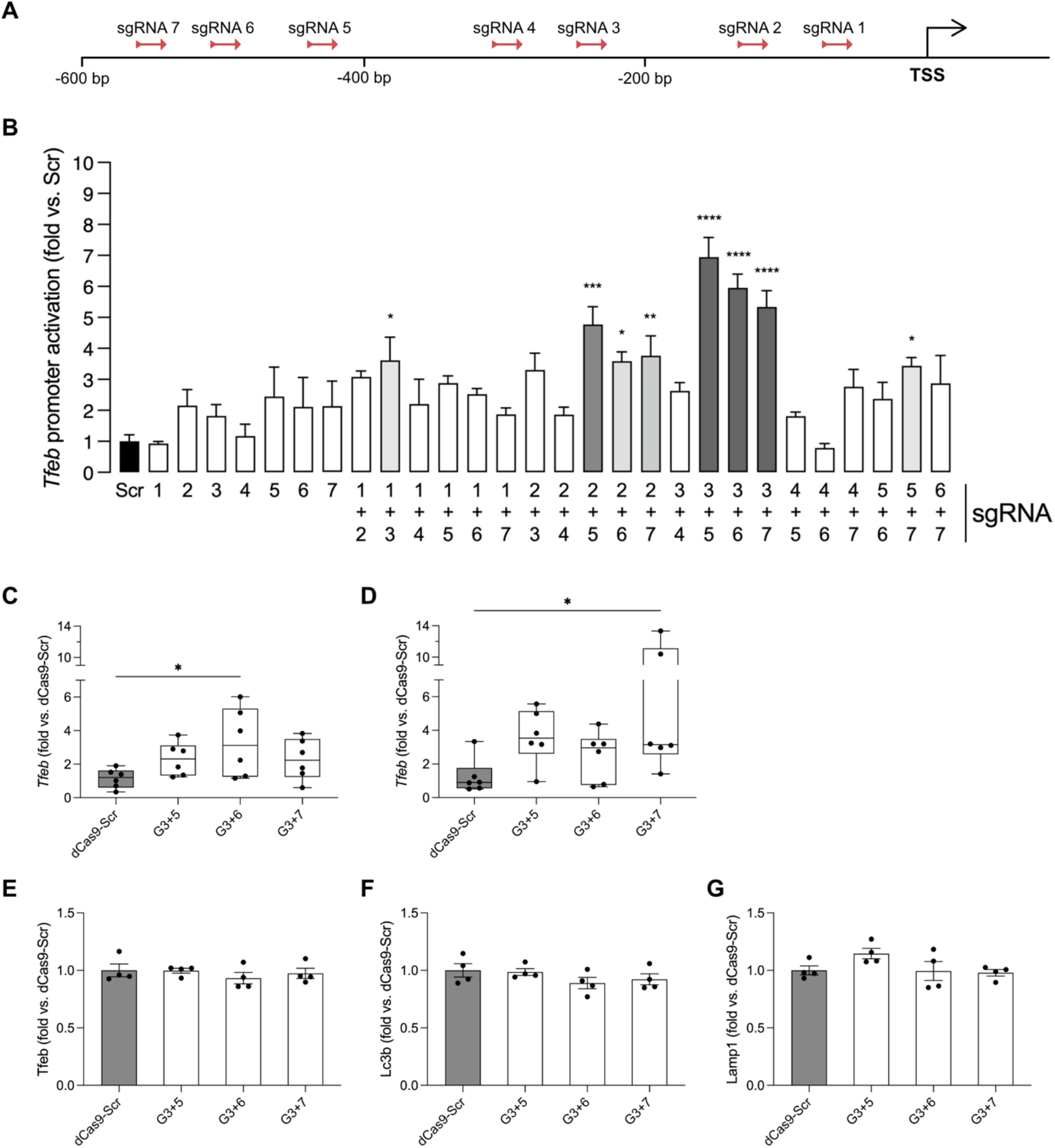
An exogenous screening assay was used to identify sgRNA candidates for experimental application. (A) sgRNA target positions relative to the transcription start site. (B) Screening results for single and co-transfected sgRNA candidates. Values are shown as fluorescence intensity relative to a non-targeting scrambled sgRNA control. All samples were normalized to a baseline BFP transfection control. RT-qPCR analysis of Tfeb expression was performed on C6 cultures (C) chemically transfected or (D) lentivirally transduced with selected sgRNA candidates. Immunocytochemistry was used to quantify (E) Tfeb, (F) Lc3b, and (G) Lamp 1 in C6 cultures lentivirally transduced with selected sgRNA candidates. (Ordinary one-way ANOVA with Dunnett’s multiple comparison test. All expression levels were calculated as fold change relative to dCas9+Scr. (*p<0.05, **p<0.01,***p<0.001, ****p<0.0001.))

RT-qPCR analysis performed on samples harvested 48 hours post-transfection revealed up to 6-fold increase in *Tfeb* expression in cultures transfected with sgRNA 3+6 compared to scrambled non-targeting control (Fig. 1C). Additionally, analysis of cultures harvested 72 hours post-LV-transduction revealed up to 13-fold increased *Tfeb* expression in samples transduced with sgRNA 3+7, albeit with notable variation in expression across replicates (Fig 1D). For all subsequent experiments, LV-mediated delivery was used. As opposed to RT-qPCR analysis, immunocytochemical (ICC) analysis of C6 cultures transduced with the respective sgRNA candidates 72 hours prior did not reveal a statistically significant increase in Tfeb protein levels (Fig. 1E). Similarly, no increase in levels of the autophagy marker Lc3b (Fig. 1F) or lysosomal marker Lamp1 (Fig. 1G) was detected, suggesting that no significant change in autophagic or lysosomal response could be determined across experimental groups.

### Assessing CRISPRa of Tfeb in a model of fibril-induced alpha-synuclein aggregation

As induced epigenetic regulation of *Tfeb* remains a poorly explored topic, we questioned whether induction of CRISPRa-mediated promoter activation could differ in a model of disease compared to a healthy cell line. Therefore, we attempted to epigenetically increase Tfeb expression in a model of α-syn aggregation.

To evaluate the effect of the CRISPRa system in a model of α-syn pathology, we first established an in vitro model of α-syn aggregation (Fig 2A). α-syn aggregation was induced through transfection of human α-syn pre-formed fibrils (PFFs) in C6 cell cultures overexpressing GFP-tagged A53T human α-syn. Bovine serum albumin (BSA) was used as a transfection control. Based on our findings in figure 1, sgRNA 3+7 was selected for LV transduction. Similar to non-transduced control cultures, C6 α-syn-GFP cultures co-transduced with sgRNA 3+7 and Mini-dCas9v2 did not display any increase in levels of Tfeb, Lc3b, or Lamp1 (Fig. 2B-D).

**Figure 2.**
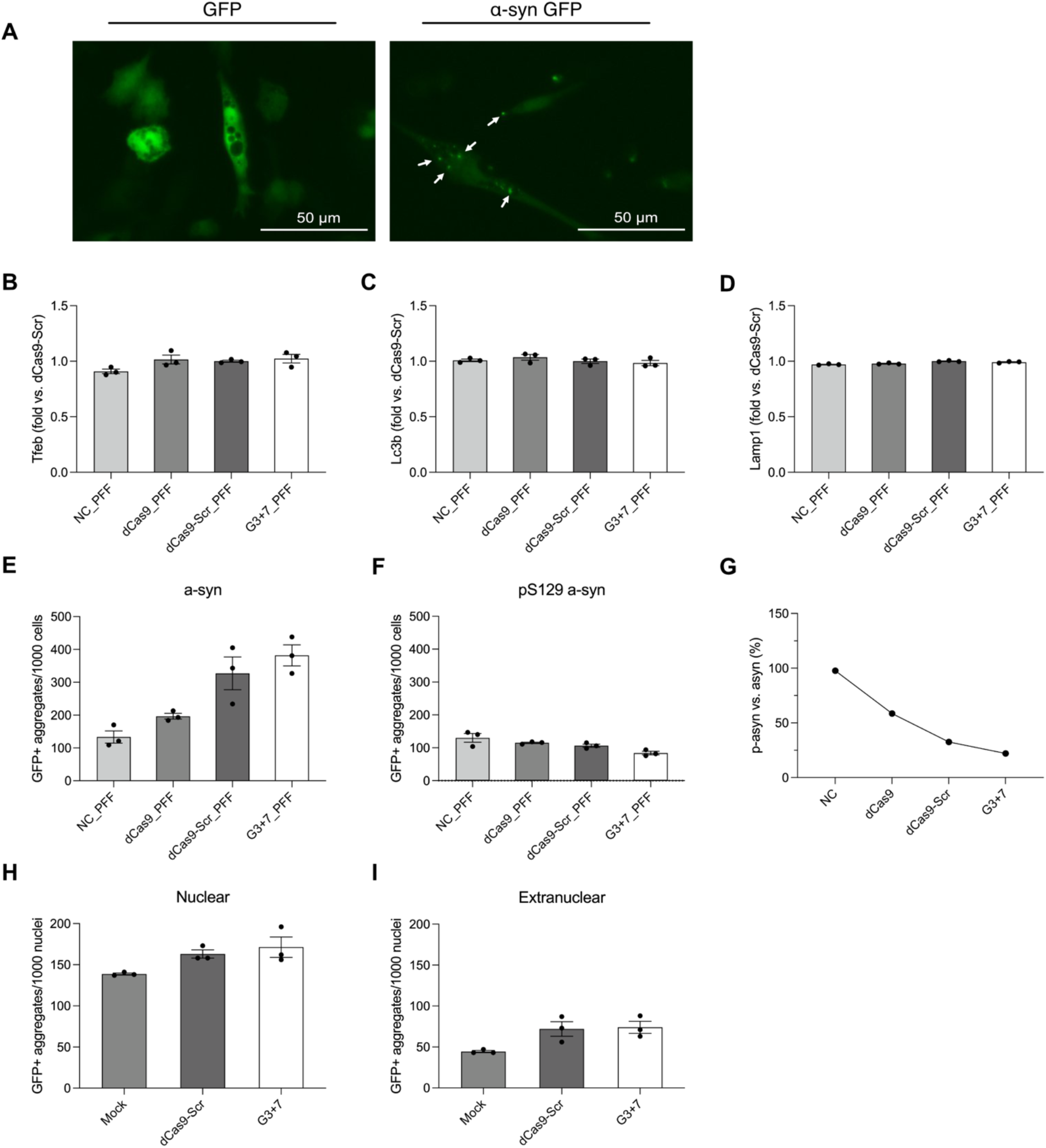
(A) An fibril seeding model of asyn-aggregation was established using C6 cells over-expressing A53T-mutated asyn-GFP. A GFP-expressing cell line was established as a control. Levels of (B) Tfeb, (C) Lc3b, and (D) Lamp1, (E) asyn, and (F) phosphorylated (pS129) asyn were quantified through immunocytochemistry in the respective conditions. (G) The relative ratio of phosphorylated-to non-phosphorylated asyn between respective conditions. (H) Nuclear GFP+ asyn aggregates detected per 1000 nuclei between respective conditions. (I) Extranuclear GFP+ asyn aggregates detected per 1000 nuclei between respective conditions.

To distinguish endogenous α-syn from transgenic α-syn-A53T-GFP overexpression, α-syn or phospho-α-syn (p-α-syn) was co-detected with a GFP-specific antibody. Administration of sgRNA 3+7 in PFF-transfected C6 α-syn-GFP cultures resulted in an increase in GFP-positive α-syn-spots compared to control settings (Fig. 2E). We did observe a stagnation in GFP-positive p-α-syn spots in the same experimental setting after transduction with sgRNA 3+7 compared to controls (Fig 2F), illustrating an over-all decreased ratio of p-α-syn to α-syn with administration of the CRISPRa system compared to controls (Fig. 2G).

We further assessed the effects on α-syn aggregation in our model by utilizing a modified version of FloIT, a previously described FACS analysis technique for selective detection of intra- and extranuclear small protein particles (26). FloIT analysis of PFF-transfected C6 α-syn-GFP cultures revealed no significant decrease in GFP-positive particle numbers in the nuclear (Fig. 2H) or extranuclear compartment (Fig. 2I). However, a slight increase in particle numbers was observed across samples treated with CRISPRa in combination with sgRNA or a non-targeting control, albeit not statistically significant.

### Combining chemical inductors of Tfeb translocation with CRISPRa

Finally, we assessed whether chemical inductions previously demonstrated to induce Tfeb activity could provide a synergistic effect with our CRISPRa system. To address this, we combined use of media containing 100 mM trehalose, 1mM valproic acid (VPA), or nutrient starvation respectively, with lentiviral transduction of the CRISPRa system in C6-cultures (Fig 3A). As VPA has been previously demonstrated to reduce viability of the C6 cell line (27), we first performed a viability assessment of C6 cell cultures treated with varying concentrations of VPA across multiple time points to determine a suitable treatment condition (Fig. S2). Based on these findings, a concentration of 1 mM VPA was selected for media supplementation.

**Figure 3.**
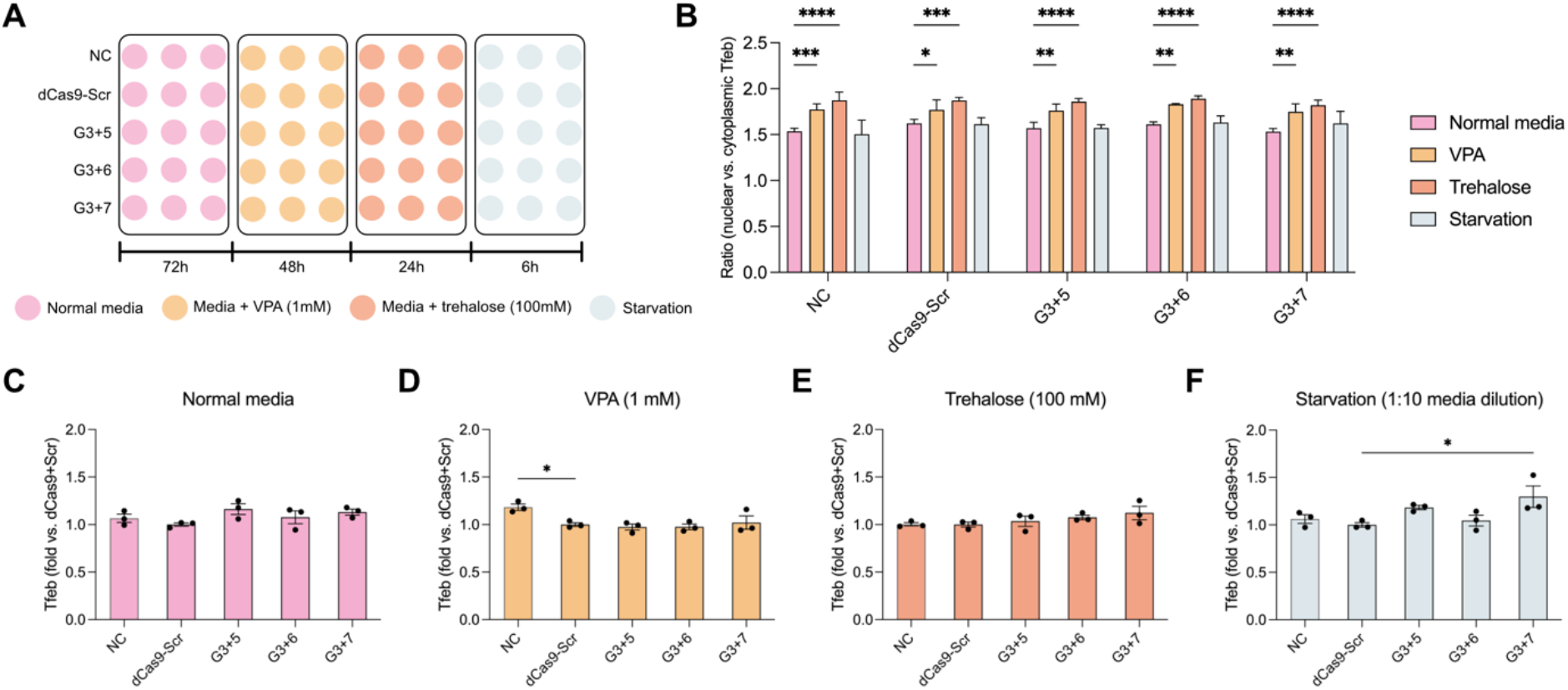
(A) Overview of the treatment timeline of LV-transduced C6 cultures. (B) Analysis of nuclear- to cytoplasmic ratio of Tfeb expression showed a general increase in nuclear Tfeb expression with VPA- or trehalose treatments across groups. No significant synergistic effect between different treatments and CRISPRa-administration was observed (Two-way ANOVA with Dunnett’s multiple comparison’s test). Nuclear Tfeb expression did not significantly increase in cultures grown in (C) normal cell culture media, (D) media supplemented with 100 mM trehalose, or (E) media supplemented with 1 mM VPA. (F) Induction of starvation response through media dilution showed an increase in nuclear Tfeb expression in cultures transduced with sgRNA 3+7. (Ordinary one-way ANOVA with Dunnett’s multiple comparison’s test). Figures C-F are expressed as fold changes relative to the average value of dCas9-Scr. (*p<0.05, **p<0.01, ***p<0.001, ****p<0.0001.))

The ratio of nuclear Tfeb expression compared to cytoplasmic levels increased across all experimental conditions upon treatment with VPA or trehalose compared to non-treated media (Fig. 3B). However, no synergistic effects of the nuclear-to-cytoplasmic ratio of expression were observed in groups transduced with the CRISPRa system compared to those transduced with a scrambled control in any of the treatment conditions (Fig. S3). When specifically analysing the levels of Tfeb present in the nucleus, a 30% increase was observed in samples transduced with sgRNA 3+7 in combination with nutrient starvation compared to the scrambled control (Fig. 3F). No other significant synergistic effects were observed (Fig. 3C-E).

## 4. Discussion

Pathological aggregation of α-syn remains a key focus in PD research. This highlights the requirement for accurate and efficient models to study accumulation of α-syn, along with experimental approaches capable of reducing build-up of toxic α-syn aggregates. In this study, we attempted to increase endogenous expression of Tfeb in healthy cell cultures and in an in vitro model of fibril-induced α-syn aggregation with the goal of reducing pathological accumulation. While we observed significant upregulation of *Tfeb* expression on mRNA level in healthy cells, we did not achieve substantial increase in protein expression in healthy or in α-syn-overexpressing cultures. Interestingly, while we observed an unexpected increase in α-syn expression across conditions, this did not correlate with an increase of phosphorylated a-syn in mock or CRISPRa treated settings.

Previous work from our group has described a streamlined workflow for cloning and screening sgRNA candidates for CRISPRa applications (24). This system utilizes a miniaturized activator complex, combining a nuclease-deficient *Staphylococcus aureus* Cas9 with a truncated VPR to achieve targeted endogenous gene activation while minimizing packaging size for downstream viral vector production (34). The sgRNA are coupled with separate hU6 promoters and packaged into cassettes flanked by compatible type-IIS (TIIS) restriction sites, allowing multiple cassettes to be assembled in tandem and incorporated in a single expression vector. This minimizes the number of constructs required for introduction of multiple sgRNA and allows for targeting of multiple sites of interest simultaneously, something that has been previously utilized particularly in CRISPRa studies (source). Furthermore, our screening method is based on plasmid transfection utilizing a construct containing the target promoter region of interest coupled to a fluorescent reporter. This allows for assessment of sgRNA binding and promoter activation independently of cell line and species and provides a middle-ground between in silico prediction and endogenous application. We have previously applied sgRNA screened this way to healthy murine cell lines and now wanted to investigate specifically *Tfeb* activation potential in a disease model.

TFEB has been a target of interest for a range of proteinopathies and accumulation disorders due to its broad involvement in autosomal-lysosomal pathway regulation. Indeed, numerous studies have demonstrated dysfunctional TFEB regulation in samples and post-mortem material from patients affected by conditions such as PD, Alzheimer’s disease, renal carcinomas, and lysosomal storage diseases (28-31). Previous work has also demonstrated a protective effect of TFEB overexpression on nigral dopaminergic neuron survival in a rat model of α-syn overexpression (16, 28). However, studies focusing on models of cardiac dysfunction have also suggested that long-term exogenous overexpression of TFEB may have harmful effects and that inducing autophagy beyond physiological levels could contribute to autophagy-dependent cell death (32, 33). We therefore opted for targeting endogenous Tfeb for upregulation, rather than employing exogenous overexpression, with the goal of sustaining a more physiologically relevant level of gene expression.

Phosphorylation of α-syn at serine 129 (pS129) has been extensively described as a marker of pathological α-syn aggregation in PD and other Lewy body diseases (35, 36). Previous work has demonstrated rapid phosphorylation of α-syn 24 hours after exposure to PFFs, illustrating the efficiency of this method of inducing aggregation in culture (37). While GFP-positive α-syn aggregates were readily detected across all PFF-seeded conditions in the FACS FloIT assay, the low levels of pS129-α-syn compared to non-phosphorylated α-syn detected may also suggest sub-par formation of phosphorylated aggregates in the C6 rat glioma cultures. Previous work has demonstrated internalization of α-syn fibrils in C6 cultures as soon as 8 hours after exposure, but did not address phosphorylation state, suggesting that pS129 α-syn induction may require further investigation in this cell model (38). For instance, a longer incubation time could allow further formation of phosphorylated aggregates. However, this approach may be unsuitable in a rapidly dividing cell line like C6.

Different forms of chemical and biological induction of TFEB activity have been extensively described, and we attempted to combine some of these with our CRISPRa system to see if we could observe synergistic effects on TFEB expression. Nutrient starvation, and particularly amino acid deprivation, naturally induces autophagy response, including TFEB translocation and activation (39). We observed increased nuclear TFEB expression upon combined nutrient deprivation and transduction of sgRNA 3+7 compared to a non-target control but could not detect a significant increase in cellular TFEB expression (Fig. S3), nor nuclear TFEB expression in scrambled control samples after nutrient deprivation compared to normal media. This suggests that our starvation model requires optimization, as nutrient deprivation-dependent nuclear translocation of TFEB has been extensively reported.

We also considered that due to the nature of TFEB as a self-regulating transcription factor, epigenetic upregulation of its expression may not automatically lead to induced autophagy response in a healthy cell setting. Trehalose is a disaccharide previously demonstrated to induce TFEB nuclear translocation, although the exact mechanism behind this has been debated (40, 41). Trehalose treatment led to an overall increased ratio of nuclear-to-cytoplasmic expression of Tfeb across our conditions, but we could not detect a significant complementary function of trehalose treatment in combination with CRISPRa.

Finally, VPA has been previously demonstrated to induce autophagy in myeloid cell models and has proven function in chromatin remodelling through inhibition of histone deacytelases (HDACs), resulting in increased chromatin accessibility and improved genome editing efficiency (42, 43). Similar to trehalose, supplementing cell media with VPA led to an overall increased ratio of nuclear-to-cytoplasmic expression of Tfeb in all experimental conditions but did not show any signs of synergistic effects with the CRISPRa system. It is worth noting, however, that we did not further assess potential changes in chromatin structure in this study Until recently, work on applied epigenetic regulation of TFEB has been sparse, and most studies have focused on translocation and quantification of the transcription factor at protein level. Naturally, this lack of literature has made it difficult to determine whether epigenetic activation of *Tfeb* is a viable, or possible, approach. However, recent work by James et al. successfully demonstrated upregulation of endogenous Tfeb in mouse osteoblast lineage cells using CRISPRa-VPR (44). To our knowledge, this is the only example of CRISPRa being successfully applied for TFEB induction with significant biological effect, confirming that induced epigenetic regulation of TFEB is a viable approach. Notably, sgRNA selected for this application were designed downstream of the transcription start site (TSS), while all sgRNA used in our study targeted the upstream region. Targeting the promoter region upstream of the TSS is typically praxis for CRISPRa applications as this provides the optimal window for induced gene expression (45, 46). Our system also includes a truncated VPR activator which has been previously characterized, but may still hold a different gene induction potential to the version used in James et al.’s work (34). Furthermore, as our system employs a dead Cas9 complex from *Staphylococcus aureus*, this imposes a more restricted PAM-targeting profile compared to SpCas9, further complicating a direct comparison between the systems and possibly limiting access to optimal target sites (47).

Our lab has previously demonstrated successful induction of endogenous *Sirt1* expression in a murine neuroblastoma cell line using the same CRISPRa system employed in this study (24). While these previous findings demonstrate the functionality of our activator system, the marked difference in gene activation between *Sirt1* and *Tfeb* raises further question regarding what makes for a suitable genetic target for this approach. A recent review by Clark and colleagues provided an extensive overview of the trials involved in selecting genomic targets, activator systems, cell- and animal models, and guide design elements to achieve activation of the target gene (48). Still, it remains difficult to say whether *Tfeb* makes a suitable candidate for CRISPRa applications, and future iterations may benefit from closer assessment of the endogenous Tfeb promoter region. This could include investigation of chromatin state, competition of the CRISPRa complex with other transcription regulators, and possibly adjusting the optimal region for sgRNA targeting beyond what is typically applied for CRISPRa. Taken together, these efforts could aid future optimization of epigenetic gene induction applications.

## Notes

### Competing Interest Statement

The authors have declared no competing interest.

## References

1. Zafar S, Lui F, Yaddanapudi SS. Parkinson Disease. StatPearls. Treasure Island (FL)2026.

2. Lanciego JL, Luquin N, Obeso JA. Functional neuroanatomy of the basal ganglia. Cold Spring Harb Perspect Med. 2012;2(12):a009621.

3. Wichmann T, Dostrovsky JO. Pathological basal ganglia activity in movement disorders. Neuroscience. 2011;198:232–44.

4. Spillantini MG, Schmidt ML, Lee VM, Trojanowski JQ, Jakes R, Goedert M. Alpha-synuclein in Lewy bodies. Nature. 1997;388(6645):839–40.

5. Stefanis L. alpha-Synuclein in Parkinson’s disease. Cold Spring Harb Perspect Med. 2012;2(2):a009399.

6. Bridi JC, Hirth F. Mechanisms of alpha-Synuclein Induced Synaptopathy in Parkinson’s Disease. Front Neurosci. 2018;12:80.

7. Wong YC, Luk K, Purtell K, Burke Nanni S, Stoessl AJ, Trudeau LE, et al. Neuronal vulnerability in Parkinson disease: Should the focus be on axons and synaptic terminals? Mov Disord. 2019;34(10):1406–22.

8. Hirsch EC, Hunot S. Neuroinflammation in Parkinson’s disease: a target for neuroprotection? Lancet Neurol. 2009;8(4):382–97.

9. Nechushtai L, Frenkel D, Pinkas-Kramarski R. Autophagy in Parkinson’s Disease. Biomolecules. 2023;13(10).

10. Kirik D, Rosenblad C, Burger C, Lundberg C, Johansen TE, Muzyczka N, et al. Parkinson-like neurodegeneration induced by targeted overexpression of alpha-synuclein in the nigrostriatal system. J Neurosci. 2002;22(7):2780–91.

11. Aniszewska A, Bergstrom J, Ingelsson M, Ekmark-Lewen S. Modeling Parkinson’s disease-related symptoms in alpha-synuclein overexpressing mice. Brain Behav. 2022;12(7):e2628.

12. Delenclos M, Burgess JD, Lamprokostopoulou A, Outeiro TF, Vekrellis K, McLean PJ. Cellular models of alpha-synuclein toxicity and aggregation. J Neurochem. 2019;150(5):566–76.

13. Volpicelli-Daley LA, Luk KC, Lee VM. Addition of exogenous alpha-synuclein preformed fibrils to primary neuronal cultures to seed recruitment of endogenous alpha-synuclein to Lewy body and Lewy neurite-like aggregates. Nat Protoc. 2014;9(9):2135–46.

14. Liu D, Guo JJ, Su JH, Svanbergsson A, Yuan L, Haikal C, et al. Differential seeding and propagating efficiency of alpha-synuclein strains generated in different conditions. Transl Neurodegener. 2021;10(1):20.

15. Decressac M, Bjorklund A. TFEB: Pathogenic role and therapeutic target in Parkinson disease. Autophagy. 2013;9(8):1244–6.

16. Arotcarena ML, Bourdenx M, Dutheil N, Thiolat ML, Doudnikoff E, Dovero S, et al. Transcription factor EB overexpression prevents neurodegeneration in experimental synucleinopathies. JCI Insight. 2019;4(16).

17. Sardiello M, Palmieri M, di Ronza A, Medina DL, Valenza M, Gennarino VA, et al. A gene network regulating lysosomal biogenesis and function. Science. 2009;325(5939):473–7.

18. Settembre C, Di Malta C, Polito VA, Garcia Arencibia M, Vetrini F, Erdin S, et al. TFEB links autophagy to lysosomal biogenesis. Science. 2011;332(6036):1429–33.

19. Liu S, Yao S, Yang H, Liu S, Wang Y. Autophagy: Regulator of cell death. Cell Death Dis. 2023;14(10):648.

20. Tan A, Prasad R, Lee C, Jho EH. Past, present, and future perspectives of transcription factor EB (TFEB): mechanisms of regulation and association with disease. Cell Death Differ. 2022;29(8):1433–49.

21. Gilbert LA, Larson MH, Morsut L, Liu Z, Brar GA, Torres SE, et al. CRISPR-mediated modular RNA-guided regulation of transcription in eukaryotes. Cell. 2013;154(2):442–51.

22. Chavez A, Scheiman J, Vora S, Pruitt BW, Tuttle M, E Pri, et al. Highly efficient Cas9-mediated transcriptional programming. Nat Methods. 2015;12(4):326–8.

23. Kunii A, Hara Y, Takenaga M, Hattori N, Fukazawa T, Ushijima T, et al. Three-Component Repurposed Technology for Enhanced Expression: Highly Accumulable Transcriptional Activators via Branched Tag Arrays. CRISPR J. 2018;1(5):337–47.

24. Arvidsson E, Lobo DD, Sabarese E, Duarte F, Nobre RJ, Quintino L, et al. A systematic screening assay identifies efficient small guide RNAs for CRISPR activation. Front Bioeng Biotechnol. 2025;13:1336313.

25. Labun K, Montague TG, Krause M, Torres Cleuren YN, Tjeldnes H, Valen E. CHOPCHOP v3: expanding the CRISPR web toolbox beyond genome editing. Nucleic Acids Res. 2019;47(W1):W171–W4.

26. Whiten DR, San Gil R, McAlary L, Yerbury JJ, Ecroyd H, Wilson MR. Rapid flow cytometric measurement of protein inclusions and nuclear trafficking. Sci Rep. 2016;6:31138.

27. Zhou Y, Xu Y, Wang H, Niu J, Hou H, Jiang Y. Histone deacetylase inhibitor, valproic acid, radiosensitizes the C6 glioma cell line in vitro. Oncol Lett. 2014;7(1):203–8.

28. Decressac M, Mattsson B, Weikop P, Lundblad M, Jakobsson J, Bjorklund A. TFEB-mediated autophagy rescues midbrain dopamine neurons from alpha-synuclein toxicity. Proc Natl Acad Sci U S A. 2013;110(19):E1817–26.

29. Kauffman EC, Ricketts CJ, Rais-Bahrami S, Yang Y, Merino MJ, Bottaro DP, et al. Molecular genetics and cellular features of TFE3 and TFEB fusion kidney cancers. Nat Rev Urol. 2014;11(8):465–75.

30. Spampanato C, Feeney E, Li L, Cardone M, Lim JA, Annunziata F, et al. Transcription factor EB (TFEB) is a new therapeutic target for Pompe disease. EMBO Mol Med. 2013;5(5):691–706.

31. Wang H, Wang R, Xu S, Lakshmana MK. Transcription Factor EB Is Selectively Reduced in the Nuclear Fractions of Alzheimer’s and Amyotrophic Lateral Sclerosis Brains. Neurosci J. 2016;2016:4732837.

32. Nah J, Zablocki D, Sadoshima J. Autosis: A New Target to Prevent Cell Death. JACC Basic Transl Sci. 2020;5(8):857–69.

33. Wundersitz S, Pablo Tortola C, Schmidt S, Oliveira Vidal R, Kny M, Hahn A, et al. The Transcription Factor EB (TFEB) Sensitizes the Heart to Chronic Pressure Overload. Int J Mol Sci. 2022;23(11).

34. Vora S, Cheng J, Xiao R, VanDusen NJ, Quintino L, Pu WT, et al. Rational design of a compact CRISPR-Cas9 activator for AAV-mediated delivery. bioRxiv. 2018:298620.

35. Anderson JP, Walker DE, Goldstein JM, de Laat R, Banducci K, Caccavello RJ, et al. Phosphorylation of Ser-129 is the dominant pathological modification of alpha-synuclein in familial and sporadic Lewy body disease. J Biol Chem. 2006;281(40):29739–52.

36. Fujiwara H, Hasegawa M, Dohmae N, Kawashima A, Masliah E, Goldberg MS, et al. alpha-Synuclein is phosphorylated in synucleinopathy lesions. Nat Cell Biol. 2002;4(2):160–4.

37. Svanbergsson A, Ek F, Martinsson I, Rodo J, Liu D, Brandi E, et al. FRET-Based Screening Identifies p38 MAPK and PKC Inhibition as Targets for Prevention of Seeded alpha-Synuclein Aggregation. Neurotherapeutics. 2021;18(3):1692–709.

38. Ihse E, Yamakado H, van Wijk XM, Lawrence R, Esko JD, Masliah E. Cellular internalization of alpha-synuclein aggregates by cell surface heparan sulfate depends on aggregate conformation and cell type. Sci Rep. 2017;7(1):9008.

39. Settembre C, Zoncu R, Medina DL, Vetrini F, Erdin S, Erdin S, et al. A lysosome-to-nucleus signalling mechanism senses and regulates the lysosome via mTOR and TFEB. EMBO J. 2012;31(5):1095–108.

40. Palmieri M, Pal R, Nelvagal HR, Lotfi P, Stinnett GR, Seymour ML, et al. mTORC1-independent TFEB activation via Akt inhibition promotes cellular clearance in neurodegenerative storage diseases. Nat Commun. 2017;8:14338.

41. Rusmini P, Cortese K, Crippa V, Cristofani R, Cicardi ME, Ferrari V, et al. Trehalose induces autophagy via lysosomal-mediated TFEB activation in models of motoneuron degeneration. Autophagy. 2019;15(4):631–51.

42. Benjamin DN, O’Donovan TR, Laursen KB, Orfali N, Cahill MR, Mongan NP, et al. All-Trans-Retinoic Acid Combined With Valproic Acid Can Promote Differentiation in Myeloid Leukemia Cells by an Autophagy Dependent Mechanism. Front Oncol. 2022;12:848517.

43. Park H, Shin J, Choi H, Cho B, Kim J. Valproic Acid Significantly Improves CRISPR/Cas9-Mediated Gene Editing. Cells. 2020;9(6).

44. James A, Hendrixson JA, Kadhim I, Marques-Carvalho A, Laster J, Crawford J, et al. Elevation of master autophagy regulator Tfeb in osteoblast lineage cells increases bone mass and strength. JCI Insight. 2025;10(17).

45. Konermann S, Brigham MD, Trevino AE, Joung J, Abudayyeh OO, Barcena C, et al. Genome-scale transcriptional activation by an engineered CRISPR-Cas9 complex. Nature. 2015;517(7536):583–8.

46. Gilbert LA, Horlbeck MA, Adamson B, Villalta JE, Chen Y, Whitehead EH, et al. Genome-Scale CRISPR-Mediated Control of Gene Repression and Activation. Cell. 2014;159(3):647–61.

47. Kleinstiver BP, Prew MS, Tsai SQ, Nguyen NT, Topkar VV, Zheng Z, et al. Broadening the targeting range of Staphylococcus aureus CRISPR-Cas9 by modifying PAM recognition. Nat Biotechnol. 2015;33(12):1293–8.

48. Clark T, Waller MA, Loo L, Moreno CL, Denes CE, Neely GG. CRISPR activation screens: navigating technologies and applications. Trends Biotechnol. 2024;42(8):1017–34.

