## Supplementary figures 1-3 for "CRISPRa-mediated activation of Tfeb in a fibril-induced alpha-synuclein aggregation model"

**Key words:**

**Supplementary material**

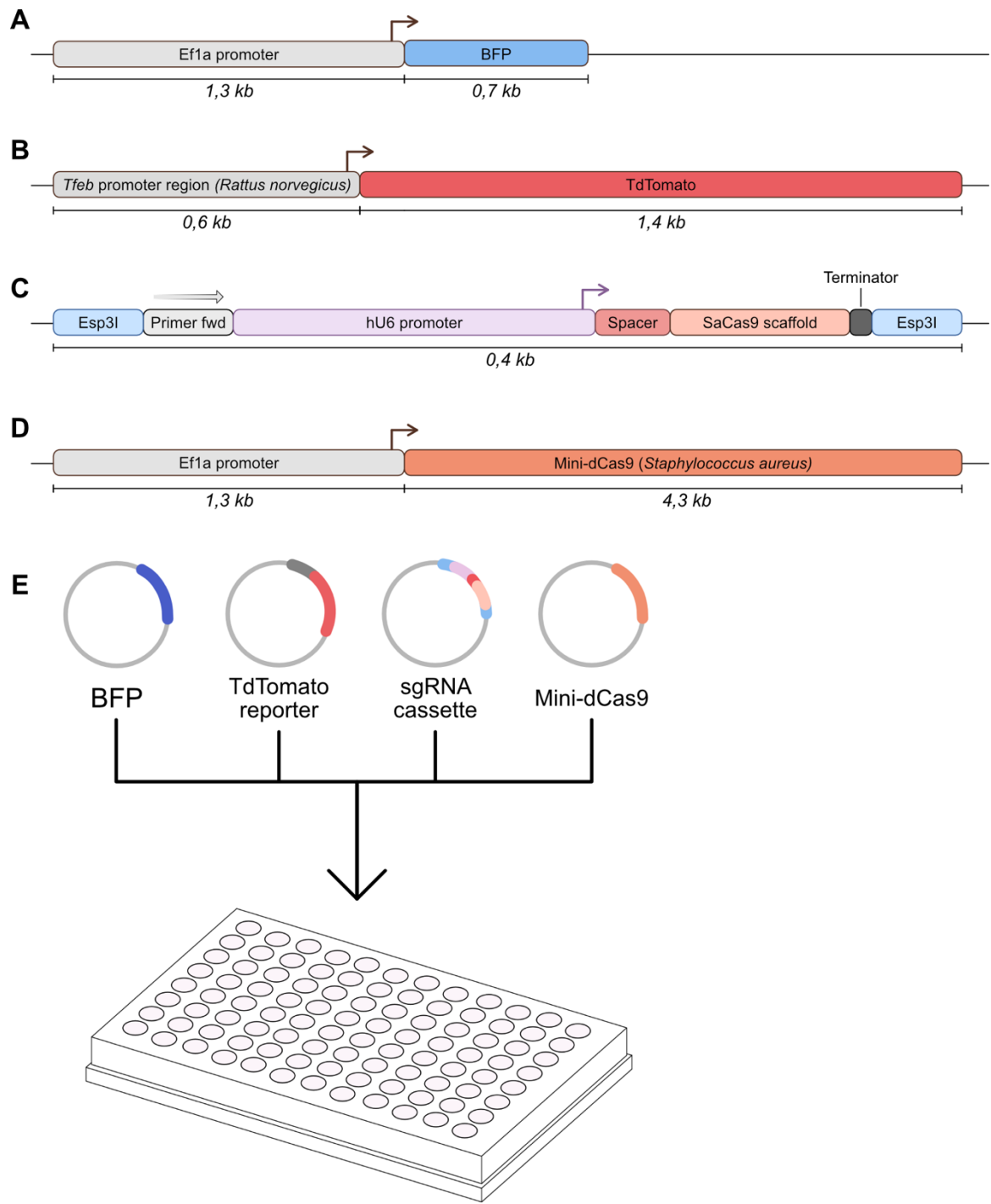

**Supplementary figure 1:** Representative plasmid maps of the (A) Ef1a-BFP transfection control, (B) TdTomato reporter, (C) the sgRNA expression cassette, and (D) the Sa-Mini-dCas9. (E) Illustration of the four-plasmid transfection for the screening assay.

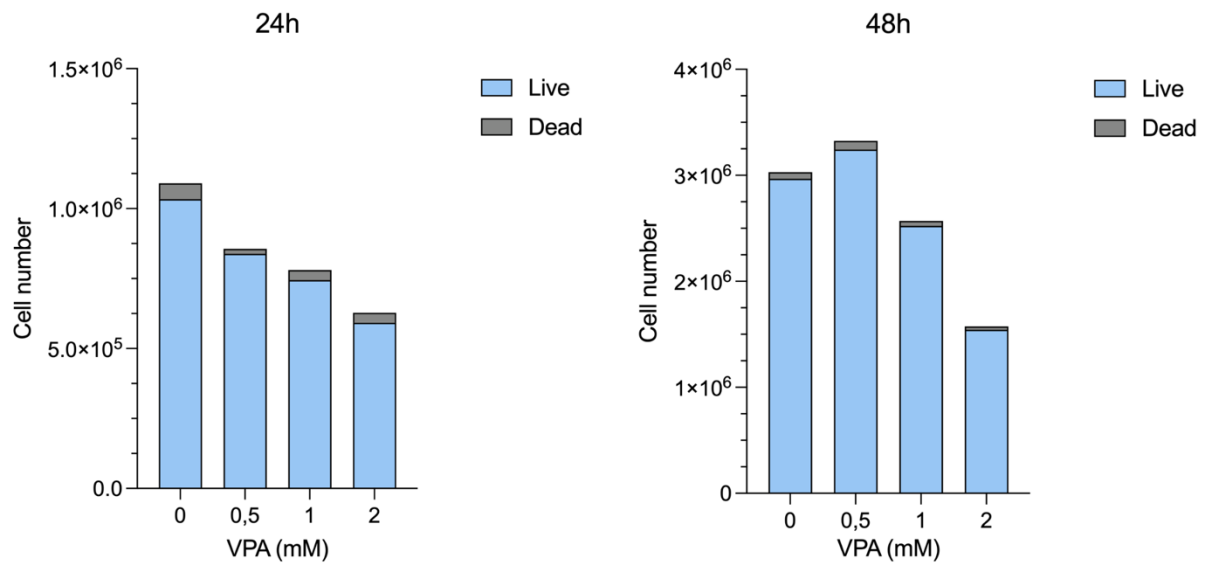

**Supplementary figure 2:** Viability assessment of C6 cultures incubated with 0-2 mM VPA over 24 and 48 hours respectively.

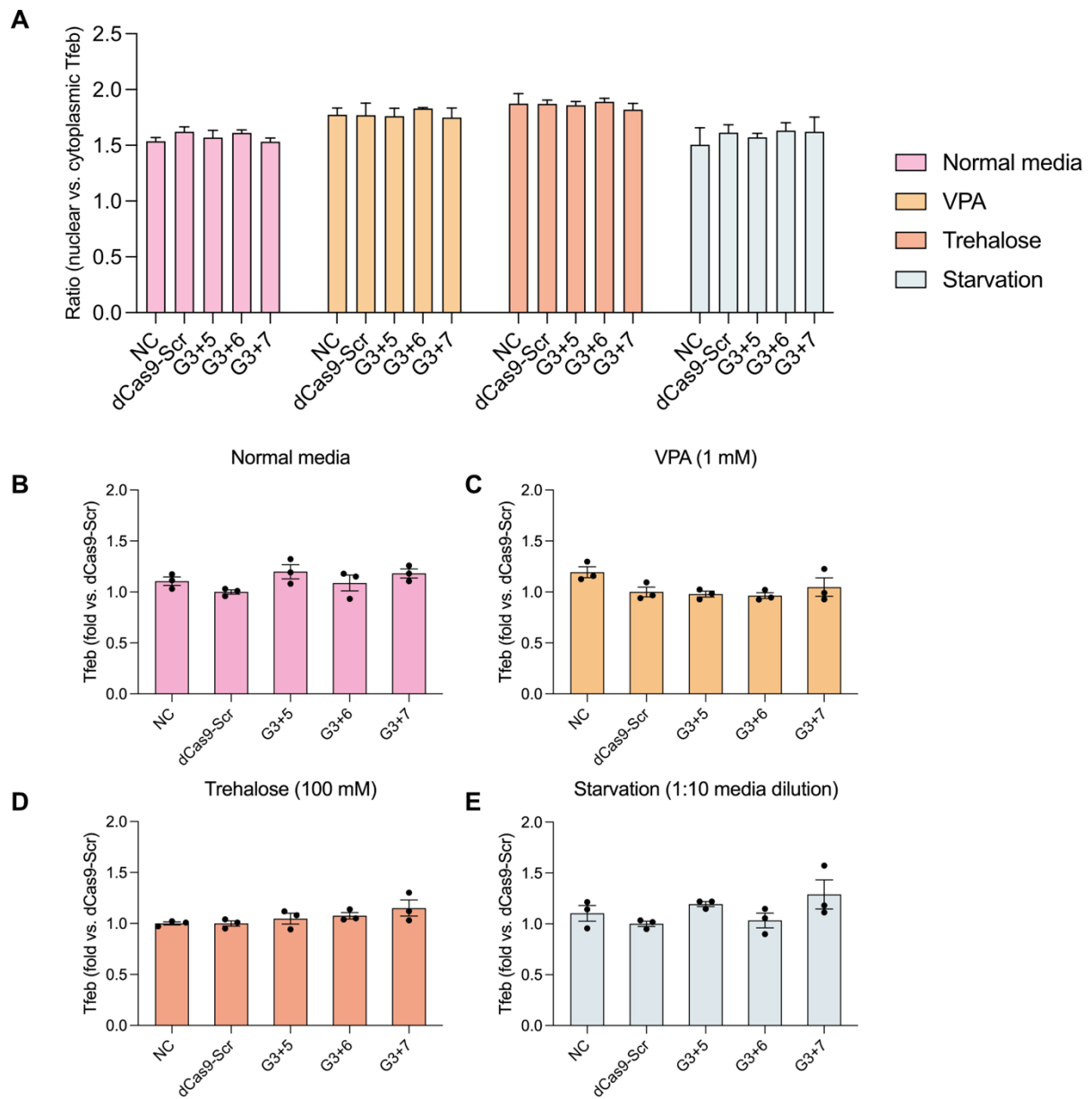

48

49 **Supplementary figure 3: (A)** Nuclear-to-cytoplasmic ratio of expression of Tfeb between media treatment
50 settings. Whole-cell expression of Tfeb in **(B)** normal media conditions, **(C)** VPA-supplemented media, **(D)**
51 **(D)** trehalose-supplemented media, and **(E)** under starvation conditions.
